# Tissue Factor Expression in Penile Squamous Cell Carcinoma: A Marker of HPV-Independent Disease

**DOI:** 10.1101/2025.06.21.658950

**Authors:** Jamaal C. Jackson, Andrew C. Johns, Leticia Campos Clemente, Christopher M. Manuel, Wei Qiao, Wei Lu, Khaja Khan, Luisa Maren Solis Soto, Jad Chahoud, Priya Rao, Matthew T. Campbell, Curtis A. Pettaway, Niki M. Zacharias

**Author notes:** Corresponding author: Niki M Zacharias. Full mailing address: Department of Urology, University of Texas MD Anderson Cancer Center, 1515 Holcombe Boulevard, Houston, TX 77030, USA. These authors contributed equally to this work (Jamaal C. Jackson, Andrew C. Johns). **Conflict of interest:** Matthew T. Campbell reports consulting and advisory roles for AXDev, Eisai, Exelixis, Merck, Pfizer, and SeaGen; institutional funding for research with Exelixis, Pfizer, Janssen, AstraZeneca, U.S. Department of Defense.

## Abstract

In a series of 33-patients, we evaluated tissue factor (TF) expression in penile squamous cell carcinoma (PSCC). A tissue microarray (TMA) was constructed with 3 cores per patient tumor (99 total cores). Anti-TF antibody staining was performed by immunohistochemistry and H-scores for membrane and cytoplasm staining were assessed (range 0-300). Percentage of cores and patient tumors staining positive for TF (≥10% of tumor cells with at least 1+ intensity in cytoplasm and/or membrane) and H-scores were described and compared with HPV and p16 status. Association of TF expression with tumor grade, presence of metastatic disease, lymphovascular invasion (LVI), perineural invasion (PNI), aberrant p53 expression, recurrence free survival (RFS), and cancer specific survival (CSS) were assessed. Nectin-4 and TROP2 staining and their association with clinical/pathological data was determined in a similar manner. TF staining was more prominent in HPV-negative tumors in both the membrane (H-score 69.6 vs 18.8; p=0.003) and cytoplasm (H-score 59.2 vs. 17.7, p=0.007). Cytoplasmic (H-score 61.7 vs 11.7, p=<0.001) and membrane TF staining (H-score 71.7 vs 15.0, p=<0.001) favored p16 negative tumors. The p53 status was more likely to be aberrant in the higher TF staining samples (cytoplasm H-score 61.7 vs 18.3, p=0.012; membrane H-score 67.5 vs 20.3, p=0.006). We observed an association with TROP2 staining and positive p16 status (membrane H-score 120.3 vs. 85, p=0.052; cytoplasmic H-score 135 vs. 107.5, p=0.041). We observed an association of TROP2 staining with positive LVI (membrane H-score 136.7 vs. 66.7, p=0.014; cytoplasmic H-score 110 vs. 93.3, p=0.04). We found no association between TF, TROP2, or nectin-4 staining with CSS or RFS; however, we suspect that this is due to our small sample size. Our results indicate that TF expression could be positive biomarker for HPV-independent, p53-aberrant PSCC, while TROP2 could be associated with HPV-associated PSCC.

## Introduction

Penile squamous cell carcinoma (PSCC) is a rare, yet aggressive malignancy with potentially devastating consequences. Patients with invasive PSCC who remain untreated typically die within 2 years due to complications from progressive locoregional or metastatic disease [1, 2]. Treatment for locoregionally advanced disease consists of either neoadjuvant cisplatin-based chemotherapy (most often in the form of paclitaxel, ifosfamide, and cisplatin (TIP)) combined with consolidative surgery. Alternatively, among patients who are not candidates for combination chemotherapy regimens, radiation-based strategies often using chemosensitization can also be used [3–7]. In patients eligible for systemic therapy, roughly 57% of patients will have evidence of disease response which yields a 4-fold improvement in median overall survival compared to non-responders when combined with surgery [8]. There are limited treatment options for patients whose cancer progresses after TIP chemotherapy, and the estimated median overall survival of non-responders ranges from less than 6 months to 17 months depending on the disease burden and whether survival was assessed from the start of treatment or at the time of clinical progression [9]. In either case the poor outcome of the chemo-refractory cohort highlights the dire need for novel therapies in this disease space.

There has been an increasing interest in exploring the use of antibody-drug conjugates (ADC) as therapeutic options in advanced PSCC. Recent analyses in PSCC have found a number of potential cellular targets, such as nectin-4 and trophoblast cell-surface antigen 2 (TROP2), that are associated with increased expression in high-risk HPV-associated PSCC when compared to HPV-negative cases [10, 11]. High expression of these molecules may be linked to contrasting clinical implications. In a recent large cohort analysis, high nectin-4 expression correlated with improved cancer-specific survival (CSS), while increased TROP2 expression was associated with early disease progression [12]. Tissue factor (TF), also known as thromboplastin or coagulation factor III, is a perivascular transmembrane glycoprotein with many functions including disruption of apoptosis via the Janus kinase (JAK)/signal transducer and activator of transcription (STAT) pathway [13]. Overexpression has been observed in numerous solid malignancies including cervical, prostate, and bladder cancer and is thought to spur primary tumor growth, neo-angiogenesis, and tumor invasion [13, 14]. The prevalence and importance to carcinogenesis make TF an intriguing potential therapeutic target. Multiple studies examining therapeutic targeting of TF in cervical cancer have shown improved efficacy and safety in the salvage setting when compared to standard therapy [15, 16]. Furthermore, the commercial availability and FDA approval of a TF-targeting ADC make it worthwhile to evaluate this target in PSCC. In this study, we evaluated TF expression in PSCC and its correlation with clinicopathological characteristics and survival outcomes.

## Methods

### Patient specimens

The current study was approved by the Institutional Review Board at The University of Texas MD Anderson Cancer Center (Protocol number 2022-0733). Patient consent was waived because the study involves no diagnostic or therapeutic intervention as well as no direct patient contact. Patients were selected from a 143-patient cohort that high-risk HPV analysis and p16 expression had previously been described by Chahoud et al [17]. These patients were further filtered to reflect those with more advanced disease by first selecting those with clinical or pathologic evidence of nodal metastasis, before eliminating patients with ≤ pathologic stage 1 disease. The cohort was ultimately selected to reflect an even distribution of HPV-associated and HPV-independent disease. Clinicopathologic data was collected including baseline characteristics, patient outcomes, and follow-up information. HPV-positive status was determined by either HPV-ISH hybridization assay or by Cobas HPV testing as described by Chahoud et al [17].

### Pathological analysis

The pathology reports in the electronic medical record were reviewed to identify appropriate achieved tissue samples obtained during the primary tumor resection. Formalin-fixed paraffin-embedded (FFPE) “donor” blocks of the selected samples were obtained. The donor blocks were then sectioned to create microscopic slides stained with hematoxylin and eosin (H&E). The H&E-stained slides were reviewed by an MDACC genitourinary pathologist (P.R.) and areas of malignant cells were marked. Once samples with sufficient tumor representative of each patient were identified, the donor blocks were used to construct a tissue microarray (TMA) with three 1.0 mm cores per tumor. A microtome was utilized to cut 5 µm sections from the cores to create TMA slides for molecular analysis [18].

### Immunohistochemistry (IHC) staining

TROP2, Nectin-4, p16, p53 and Human Tissue Factor-1 (CD142) markers have been studied with IHC assays. Four-micron thickness formalin-fixed paraffin embedded (FFPE) tissue sections were mounted on positively charged glass slides. The section went through standard staining protocols on either Leica BOND™ Autostainer (Leica Biosystems, USA) with BOND™ Polymer Refine Detection kit (Leica Biosciences, DS9800), or Ventana Discovery Ultra (Ventana Medical System, USA) with OptiView DAB Detection Kit (Roche, 760-700). The antibody clone, catalogue number, vendor, dilution factor, antigen retrieval condition, antibody incubation time, and staining platform are summarized in **Supplemental Table 1**. Briefly, the tissue sections were baked and deparaffinized, followed by epitope retrieval, then hydrogen peroxide blocking, then incubated with primary antibody and detection system. The antibody-antigen binding was visualized by brown DAB Refine stain and counter-stained blue with Hematoxylin for 8 minutes. Tissue slides were then dehydrated offline, cleared in xylene, and cover-slipped using Cytoseal™ XYL solution. The stained slides were scanned with Leica Aperio AT2 slide scanner for analysis.

### Immunohistochemistry (IHC) scoring methodology and analysis

Anti-TF-1 antibody staining was performed by immunohistochemistry and evaluated in the membrane and cytoplasm of malignant cells and reported as histologic score (H-score) considering the extension (percentage) of positive expression at each intensity level (0, no staining, 1+ mild staining, 2+ moderate staining, 3+ strong staining) (range 0-300). H-scoring was also evaluated for nectin-4 and TROP2 membranous and cytoplasmic expression in malignant cells. Tumor protein p53 immunohistochemistry interpretation was performed based on the criteria initially outlined for vulvar cancer [19, 20] extended to penile cancer [21]. Normal p53 expression was characterized by either patchy nuclear staining with variable intensity or moderate to strong nuclear staining sparing the basal layer. Abnormal p53 expression was defined by one of four distinct patterns: (1) continuous strong and nuclear staining restricted to the basal layer, (2) diffuse strong nuclear staining least of at least 80% of cells, (3) complete absence of nuclear staining in all tumor cells, with evidence of positive internal control, or (4) moderate to strong cytoplasmic staining. The p16 immunohistochemistry was evaluated based on the hybrid system (HS) previously described by Chahoud et al [17]. Tumor sections were considered positive if one of the following staining patterns was present in more than 75% of neoplastic cells (1) extensive discontinuous or (2) entire and continuous cytoplasmic or nuclear. Percentage of tumors staining positive for TF, TROP2, and nectin-4 (≥ 10% of tumor cells with at least 1+ intensity in cytoplasm and/or membrane) and median H-scores were described and compared based on specimen HPV status, p53, and p16 staining.

### Study endpoints and follow-up

Patients were followed from the time of definitive surgical intervention until the last recorded assessment in the medical record or death. Patient were grouped into one of 4 categories at the time of last follow-up: “Alive without disease” “Alive with disease”, “Dead of disease”, and “Deceased from another cause”. Tumor specimen grade was classified using the American Joint Committee on Cancer (AJCC) 8^th^ edition staging system criteria. Association of TF, TROP2, and nectin-4 expression with tumor grade, presence of metastatic disease, p53 staining pattern, p16 expression, lymphovascular invasion (LVI), perineural invasion (PNI), recurrence free survival (RFS), and cancer specific survival (CSS) was assessed. RFS was calculated as time from the date of surgery to the first recurrence (either local, regional, or distant) or death or censored at the last follow-up. CSS was calculated as time from the date of diagnosis to cancer related death or censored at the last follow-up.

### Statistical analysis

Numerical variables are reported using the median, minimum, and maximum value. Categorical variables are reported with frequencies and percentages. Fisher’s exact test was used to assess association between categorical variables and TF, TROP2, and nectin-4 average staining scores, while the Wilcoxon rank sum test was used to compare numerical variables which were expressed with median and interquartile range (IQR). Results for all associations determined using categorical variables are reported in **Supplemental Table 2**. Spearman’s method was calculated to assess the degree of correlation (denoted as r) between the membrane and cytoplasm specific average staining scores, respectively. Cox Proportional Hazard models were constructed to assess the association between the different H-scores, HPV, p16 and aberrant p53 status with RFS and CSS. For all tests, a α=0.05 is used as the threshold for statistical significance. All computations were carried out using R version 4.4.1.

## Results

### Patient baseline characteristics

This study included 33 patients who underwent primary surgical treatment of PSCC at MDACC. Patient demographics for our cohort are provided in **Supplemental Table 3**. The majority of these patients were Caucasian (60.6%) and underwent partial penectomy as their primary treatment (66.7%). Median age was 64 years (IQR 51.5 – 70). Pertinent patient history included significant tobacco use (60.6%), phimosis (57.6%), and lack of neonatal circumcision (12.1%). There was general concordance on tumor assessment as most lesions were categorized as clinical stage T2 (51.5%) and N0 (57.6%) versus pathologic stage T2 (69.7%) and N0 (36.4%) on pre- and post-surgical analysis, respectively. Lymphovascular and perineural invasion were noted in 24 (72.7%) and 12 (36.4%) patients, respectively. Seventeen patients (51.5%) had HPV-positive disease while 19 patients (57.6%) had metastatic disease. Patients were followed for a median of 19.3 months (IQR 5.5 – 42.2 months). Most patients were classified as “alive without disease” (42.4%) at the time of this study.

### Study endpoint results

Positive staining for tissue factor (TF) was detected in 27 of 32 evaluable tumors (84.4%). Of the tumors that failed to stain, 4 out of the 5 were HPV-positive (80%). Median overall membrane TF H-score was 47 (range 1-232), while median cytoplasmic H-score was slightly lower at 29 (range 2 – 180). All 32 tumors stained positive for TROP2. The TROP2 median membrane H-score was 106 (range 0-277) and the median cytoplasmic H-score was 109 (range 40-227). The lowest percentage of positive staining was seen with nectin-4. Only 15 of the 32 tumors (47%) were positive for nectin-4 by membrane staining, and the median H-score was 5.4 (range 0-137). Twenty-four of the 32 tumors (75%) were positive for nectin-4 by cytoplasmic staining, and the median H-score was 74 (range 0-172). Representative images of 3 tumor tissues with variable expression in TF, TROP2, and nectin-4 are shown in **Figure 1**.

**Figure 1.**
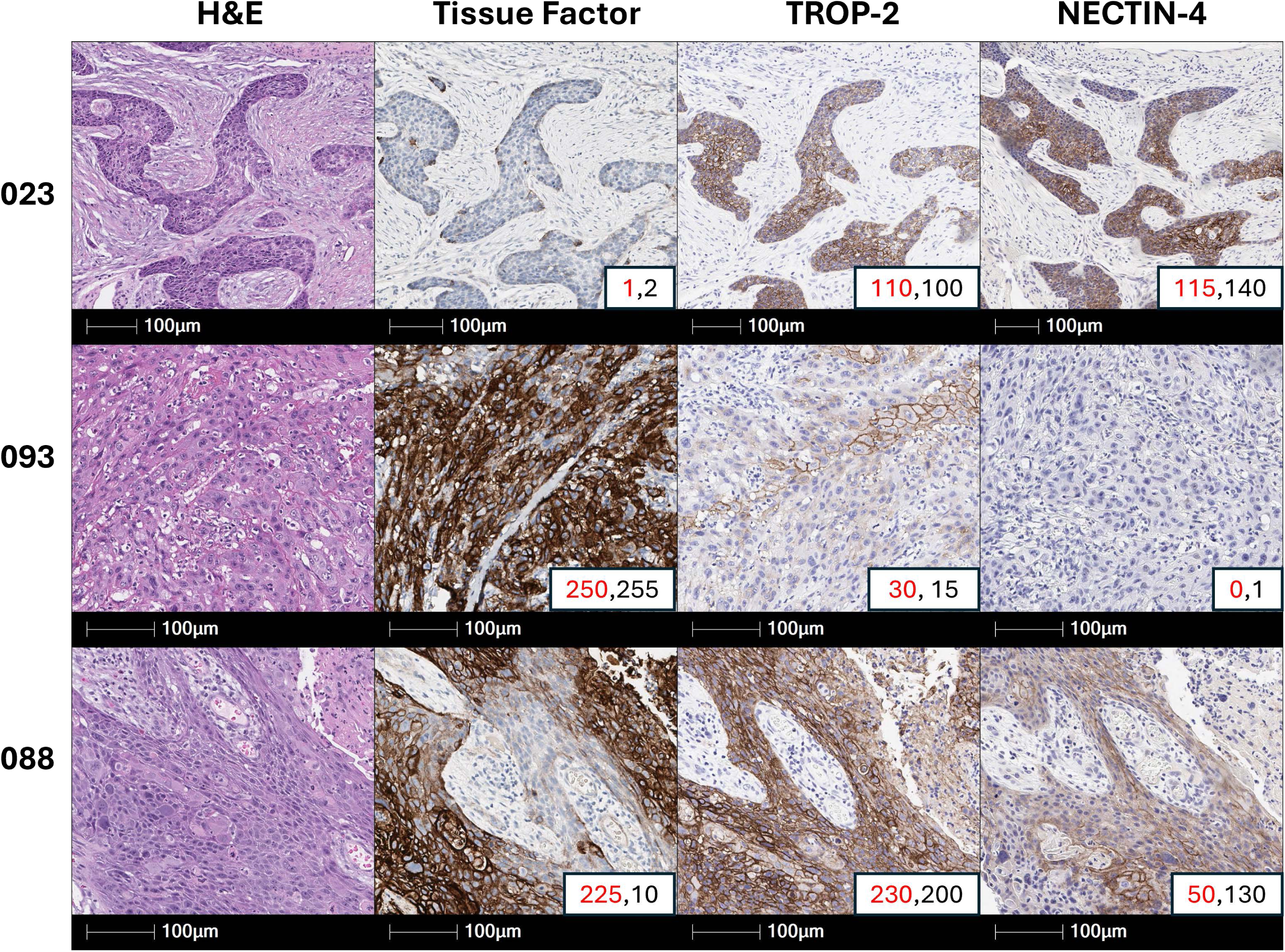
Representative images of three tumor tissues with variable expression of tissue factor (TF), TROP2, and nectin-4. Membrane H-score is in red font the cytoplasmic H-score is in black font in lower-right hand corner of each image.

### Associations with TF expression

A significant association was observed between HPV-status and TF expression. Staining was more prominent in HPV-negative tumors in both the membrane (median H-score 69.6 vs 18.8; p=0.003) and cytoplasm (median H-score 59.2 vs. 17.7, p=0.007). A similar result was observed when comparing TF expression based on p16 status. Cytoplasmic (61.7 vs 11.7, p=<0.001) and membrane TF staining (71.7 vs 15.0, p=<0.001) favored p16 negative tumors. Final p53 status was more likely to be aberrant in the higher staining samples (cytoplasm: 61.7 vs 18.3, p=0.012; membrane: 67.5 vs 20.3, p=0.006). There was no association with either membrane [HR 1.0 (95% CI 0.99 – 1.01), p=0.896] nor cytoplasmic TF staining [HR 1 (95% CI 0.99 – 1.01), p=0.729] when assessing CSS. Similarly, no association was observed when using RFS as an endpoint in both membrane [HR 1 (95% CI 0.99 – 1.01), p=0.535] or cytoplasmic TF staining [HR 1 (95% CI 0.99 – 1.01), p=0.514]. There were no significant associations linking TF H-score to PNI, LVI, metastasis, or primary tumor grade.

### Associations with TROP2/Nectin-4 Expression

Unlike TF, we did not observe any association with TROP2 and nectin-4 staining with aberrant p53 or HPV status. We observed an association of TROP2 staining with positive LVI (membrane median H-score 136.7 vs. 66.7, p=0.014; cytoplasmic median H-score 110 vs. 93.3, p=0.04). We observed an association with TROP2 staining and positive p16 status (membrane 120.3 vs. 85, p=0.052; cytoplasmic 135 vs. 107.5, p=0.041). Similar to TF, we did not observe any correlations with RFS and CSS with TROP-2 or nectin-4 staining.

### Correlations Between Surface Proteins and Other Factors

We estimated the spearman correlation between TF, nectin-4, and TROP2 using the membrane staining scores (**Figure 2A**) and cytoplasmic staining scores (**Figure 2B**). Correlations that are off the diagonal are statistically significant. We observed a positive correlation between TROP2 and nectin-4 membrane staining (r = 0.41), while we observed a negative correlation between cytoplasmic TF and TROP2 staining (r = −0.42). Contingency tables were utilized to determine the association between aberrant p53 and HPV status; aberrant p53 and p16 status; and p16 and HPV status (**Table 1**). The p-values obtained from the Fisher’s exact test confirmed that the three sets of variables were all associated with each other. In addition, we determined the association of p16, HPV, and aberrant p53 status with CSS and RFS. **Table 2** shows these associations along with hazard ratios (HR). Positive HPV and p16 status are marginally associated with RFS at an α = 0.10, both exhibiting a protective effect on recurrence. Aberrant p53 is associated with RFS (p-value = 0.007) and has a negative effect on RFS. For CSS, positive HPV is marginally associated, while positive p16 status is associated with a p-value = 0.024. As with RFS, positive HPV and p16 status exhibit a protective effective for CSS. For p53 and CSS, the large HR coefficient is probably due to the low number of cancer related deaths with normal p53 staining (only 1 death).

**Figure 2.**
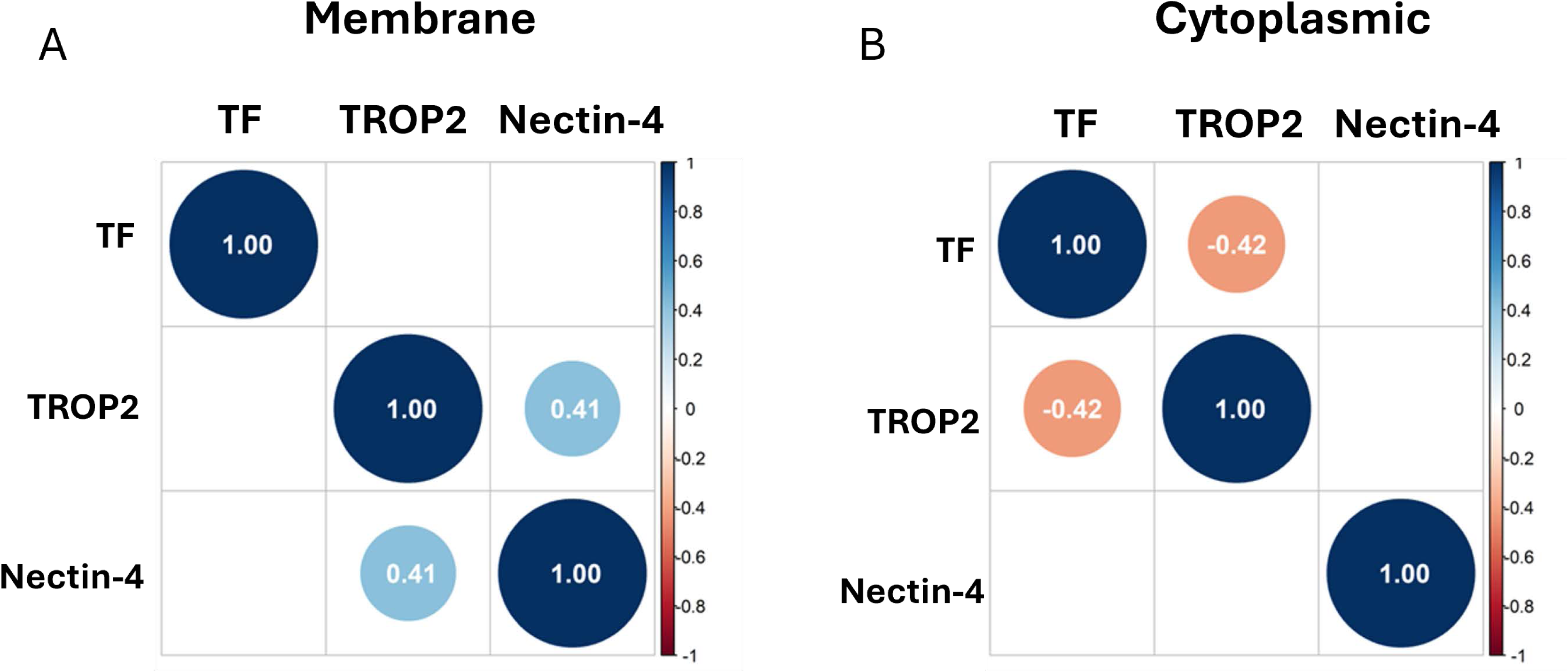
Correlation Matrices. Correlations that are off the diagonal are statistically significant, with positive correlations shown in light blue circles and negative correlations shown in orange circles. Correlation plot of the membrane staining (A) and of the cytoplasmic staining (B) of TF, TROP2, and nectin-4 shown.

**Table 1.** Contingency tables illustrating the associations between aberrant p53 and HPV status; aberrant p53 and p16 status; and p16 and HPV status.

| Patient characteristics | Total | HPV-negative | HPV-positive | p-value |
| --- | --- | --- | --- | --- |
| p53 status |  |  |  |  |
| normal | 21 (65.6%) | 7 (43.8%) | 14 (87.5) | 0.023 |
| aberrant | 11 (34.4%) | 9 (56.2%) | 2 (12.5%) |  |
| p16 status |  |  |  |  |
| negative | 17 (53.1%) | 15 (93.8%) | 2 (12.5%) | < 0.001 |
| positive | 15 (46.9%) | 1 (6.3 %) | 14 (87.5%) |  |
| Patient characteristics | Total | p16 negative | p16 positive | p-value |
| p53 status |  |  |  |  |
| normal | 21 (65.6%) | 6 (35.3%) | 15 (100) | < 0.001 |
| aberrant | 11 (34.4%) | 11 (64.7%) | 0 (0%) |  |

**Table 2.** Associations with RFS and CSS with HR for HPV, p16, and aberrant p53 status.

| RFS |  |  |  |
| --- | --- | --- | --- |
| Variable | Level | HR (95% CI for HR) | p-value |
| HPV | positive vs. negative | 0.45 (0.17-1.17) | 0.103 |
| p16 | positive vs. negative | 0.4 (0.15-1.07) | 0.069 |
| p53 | aberrant vs. normal | 4.06 (1.48-11.15) | 0.007 |
| CSS |  |  |  |
| Variable | Level | HR (95% CI for HR) | p-value |
| HPV | positive vs. negative | 0.23 (0.04-1.2) | 0.081 |
| p16 | positive vs. negative | 0.09 (0.01-0.73) | 0.024 |
| p53 | aberrant vs. normal | 4683379200 (0-infinity) | 0.999 |

## Discussion

This study examined a cohort of PSCC patients with well-documented clinical outcomes to assess TF expression in primary tumor specimens. We found significant TF positivity in 84.4% of the cases using a cut off of ≥ 10% of tumor cells with at least 1+ intensity in cytoplasm and/or membrane. When examined in the context of HPV-status, this encompassed 93.8% positivity in HPV-negative cases and 75% in HPV-positive specimens. We found TF expression was highly associated with p16 and HPV-negative tumors and aberrant p53 staining. Similar results were not observed with TROP2 and nectin-4 staining. In addition, a negative correlation was observed with membrane TF and TROP2 staining, indicating that these two surface proteins could be expressed in two different subsets of patients – TROP2 in HPV-positive tissues and TF in HPV-negative tissues. Our results indicate that TF expression could be positive biomarker for HPV-independent PSCC, which is currently determined by the lack of p16 staining and a negative test for HPV using either PCR or *in situ* hybridization assays. HPV-independent PSCC has been shown to be associated with *TP53* mutations [21–23]. The results from contingency Table 1 validates this assumption as we see aberrant p53 is associated with negative p16 and HPV status. Our TF data suggests that TF might be a driver/passenger in *TP53* mutated HPV-independent PSCC.

Our TF values are on par with the reported rates of TF expression in other solid tumors highlighted in a recent study by de Bono et al [24]. Criteria for positive TF expression in that study were similar to our study and were characterized by modified H-scores. It should be noted that a number of tumors analyzed in the study by de Bono et al utilized tissue obtained from biopsy as opposed to completely excised tissue which may have impacted the observed positivity rates. Interestingly, cervical cancer, which has similar pathogenesis to PSCC, showed TF expression of 77%. Other studies have also corroborated this increased expression in cervical cancer and described an association of increased expression with worse outcomes [25]. It is important to note that unlike PSCC, which is only associated with high-risk HPV infection in up to 50% of cases, cervical cancer is almost universally associated with infection with high-risk HPV [26].

To our knowledge, this is the only study to date highlighting increased TF expression in HPV-negative PSCC specifically. The majority of studies exploring novel targets in PSCC have largely focused on HPV-positive PSCC. Grass et al. first described elevated expression of nectin-4, an immunoglobulin involved in cell-cell junction formation and maintenance, in invasive PSCC in 2022 [11]. Programmed death ligand-1 (PD-L1) and TROP2 have also been examined with associations found between these proteins and higher tumor stage and disease progression, respectively [12]. Prior studies have noted increased expression of both TROP2 and nectin-4 in HPV-positive cases, but targets for HPV-negative PSCC are sparse in the literature. In our study, we did not find a significant association of TF H-score with tumor grade, presence of metastatic disease, LVI, PNI, CSS, or RFS. Previous literature has detailed the association of TF expression with poor prognosis in other solid tumors [27]. Associations between TF overexpression and metastases in breast cancer, non-small cell lung cancer, and colorectal cancer have been reported [28–30]. Furthermore, increased TF expression in colorectal cancer has specifically been linked to the development of hepatic metastasis, which carries a particularly grim prognosis [31]. TF was not associated with RFS, CSS, or metastatic progression in our study, however this may be a product of the small sample size. The limitations of a small sample size in our study are evident in the large, calculated HR for aberrant p53 status in CSS, as it is driven by the fact that only one cancer-related death occurred in the p53-negative cohort. However, even in this small cohort aberrant p53 was associated with worse RFS (Table 2). Our data is consistent with a recent report from Elst et al revealing that aberrant p53 tumor expression was associated with worse CSS among a cohort of 541 subjects irrespective of HPV status [32]. Given the TF identifies the aberrant p53 cohort and represents a viable therapeutic target, the current study is particularly relevant and warrants further investigation of TF expression and its association with PSCC progression in a larger patient cohort.

As previously stated, prognosis in advanced PSCC remains poor in those whose disease has progressed on frontline platinum-based systemic therapy. These outcomes are decidedly worse in those who present with HPV-independent disease. The findings of increased TF expression in this population may present an ideal therapeutic target as treatment in similar tumors has yielded notable results. For example, cervical cancer has also been demonstrated to have TF expression of up to 77% which exceeds expression in other solid tumors [24]. Tisotumab vedotin (TV) is a TF-specific antibody-drug conjugate paired with monomethyl auristatin E (MMAE) that works to inhibit microtubule activity; thus, enacting a direct cytotoxic effect [33]. Coleman et al. conducted a multicenter phase II trial utilizing TV in patients with recurrent or metastatic cervical cancer who had progressed on or after doublet chemotherapy with bevacizumab and had two or fewer previous systemic regimens for their disease in the frontline setting. This study showed a confirmed objective response rate of 24% with 28% of study patients reporting grade 3 or worse treatment-related adverse events (TRAEs) [15]. A follow-up phase 3 study examining TV as second- or third-line therapy in patients with recurrent or metastatic cervical cancer demonstrated a similar improvement in confirmed objective response rate and TRAEs, in addition to a 30% lower risk of death and improved progression-free survival in the TV cohort when compared to standard chemotherapy regimens [16]. These results support the safety of TV and other ADCs in the clinical setting and its potential for use in PSCC in the second-line setting and beyond. In locally advanced cases, patients who show a significant response may also be more likely to benefit from consolidative therapies (e.g. radical surgical resection or radiation).

There are several limitations of this study. Firstly, the sample size of only 33 patients is relatively small, but consistent with the rarity of PSCC. Secondly, the most recent primary surgery from the study occurred in 2016 which makes all samples 8+ years old at the time of the analysis. This may introduce time-based degradation of the antigens in question and may have impacted the fidelity of immunohistochemical staining and scoring. Lastly, utilizing TMAs in our analysis may not have fully accounted for tumor heterogeneity as this represents a small sample of the overall malignancy, in contrast to examination of whole tissue slides.

The role of TF in PSCC is a promising development that may have important treatment implications in the future. Preclinical studies will be of interest to further delineate the benefit of targeting TF in PSCC with eventual transition to clinical trials in this space. This is already being explored in other ADCs with great anticipation [10]. With continued research, one can foresee incorporation of targeted therapy into the PSCC treatment paradigm in the future and hopefully improvement in clinical outcomes.

## Conclusion

TF expression was noted among the majority of PSCC with expression at rates comparable to published data for cervical cancer. Expression was significantly higher among the HPV and p16 negative cohort and the aberrant p53 cohort. These data support further investigation of TF-targeted therapy, such as an ADC, for treatment of HPV-independent PSCC.

### Ethics approval

This study was approved by the Institutional Review Board at The University of Texas MD Anderson Cancer Center (Protocol number 2022-0733). Patient consent was waived because the study involves no diagnostic or therapeutic intervention as well as no direct patient contact.

### Author Contributions

A.C.J, J.C.J, N.M.Z, M.C., L.S.S., and C.A.P conceived the project. N.M.Z and J.C.J pulled the blocks and cut pathology slides. N.M.Z. and C.A.P. provided funding. P.R. reviewed the H&E slides. L.C.C. and L.S.S. scored all IHC staining. W.L. and K.K. performed TF staining. A.C.J. and J.C.J. reviewed all clinical data. C.M.M. and W.Q. performed all statistical analysis. A.C.J., J.C.J., C.A.P., N.M.Z., L.C.C., L.S.S., C.M.M., and W.Q. wrote initial manuscript. All authors edited and approved the final manuscript.

## Data availability

The data presented in this study are available upon reasonable request from the corresponding author.

---

**Supplemental Table 1.** The antibody clone, catalogue number, vendor, dilution factor, antigen retrieval condition, antibody incubation time, and staining platform are summarized.

| **Antibody** | **Clone** | **Vendor** | **Cat#** | **Dilution Factor** | **Antigen Retrieval** | **Antibody Incubation Time** | **AutoStainer** |
| --- | --- | --- | --- | --- | --- | --- | --- |
| **TROP2** | EPR20043m | Abcam | ab214486 | 1:4000 | ER1 for 20 minutes at 100°C | 15 minutes | Leica Bond III platform |
| **NECTIN-4** | EPR15613-68 | Abcam | ab192033 | 1:1000 | ER2 for 10 minutes at 100°C | 15 minutes | Leica Bond III platform |
| **P16** | E6H4 | Ventana | 705-4793 | Ready to Use | CC1 for 48 minutes at 100°C | 32 minutes | Benchmark Ultra Platform |
| **P53** | DO-7 | Leica | PA0057 | 1:2 | ER2 for 20 minutes at 100°C | 15 minutes | Leica Bond III platform |
| **CD142** | HTF-1 | Thermo Fisher | #16-1429-82 | 1:200 | CC1 for 80 minutes at 100°C | 32 minutes | Discovery Ultra Platform |

Note: ER1: Epitope Retrieval solution 1, pH 6.0; ER2: Epitope Retrieval solution 2, pH9.0; CC1: Cell condition Ultra, pH8.0.

**Supplemental Table 2.** Associations determined using average H-score of TF, TROP2, and nectin-4 with PNI, LVI, p53, HPV, primary tumor grade, and p16.

PNI
| Patient Characteristics |  | Total N=32 | no N=10 | yes N=12 | P-value |
| --- | --- | --- | --- | --- | --- |
| Membrane Tissue Factor Average Final H Score, median (min,max) | N = 32 | 46.92 (0.67, 231.67) | 69.59 (5.67, 136.67) | 43.75 (4, 231.67) | 0.722 |
| Cytoplasm Tissue Factor Average Final H Score, median (min,max) | N = 32 | 29.16 (2, 180) | 33.5 (5, 98.33) | 31.09 (3, 180) | 0.921 |
| Trop 2 Membrane Average Final H Score, median (min,max) | N = 32 | 106.66 (0, 276.67) | 111.66 (66.67, 276.67) | 98.34 (0, 276.67) | 0.276 |
| Trop 2 Cytoplasm Average Final H Score, median (min,max) | N = 32 | 109.16 (40, 226.67) | 111.66 (93.33, 226.67) | 100 (40, 210) | 0.106 |
| Nectin 4 Membrane Average Final H Score, median (min,max) | N = 32 | 5.42 (0, 136.67) | 10.16 (0, 69.33) | 3.33 (0, 115) | 0.712 |
| Nectin 4 Cytoplasm Average Final H Score, median (min,max) | N = 32 | 74.16 (0, 171.67) | 90 (3.33, 166.67) | 58.34 (0, 168.33) | 0.337 |

LVI
| Patient Characteristics |  | Total N=32 | no N=7 | yes N=23 | P-value |
| --- | --- | --- | --- | --- | --- |
| Membrane Tissue Factor Average Final H Score, median (min,max) | N = 32 | 46.92 (0.67, 231.67) | 47.5 (17.5, 231.67) | 46.33 (1.33, 205) | 0.471 |
| Cytoplasm Tissue Factor Average Final H Score, median (min,max) | N = 32 | 29.16 (2, 180) | 30.5 (18.33, 180) | 28.33 (2.33, 123.33) | 0.202 |
| Trop 2 Membrane Average Final H Score, median (min,max) | N = 32 | 106.66 (0, 276.67) | 66.67 (0, 105) | 136.67 (15, 276.67) | 0.014 |
| Trop 2 Cytoplasm Average Final H Score, median (min,max) | N = 32 | 109.16 (40, 226.67) | 93.33 (40, 136.67) | 110 (73, 226.67) | 0.041 |
| Nectin 4 Membrane Average Final H Score, median (min,max) | N = 32 | 5.42 (0, 136.67) | 0 (0, 69.33) | 17 (0, 136.67) | 0.117 |
| Nectin 4 Cytoplasm Average Final H Score, median (min,max) | N = 32 | 74.16 (0, 171.67) | 46.67 (0, 90) | 90 (0, 171.67) | 0.073 |

p53 Status
| Patient Characteristics |  | Total N=32 | normal N=21 | aberrant N=11 | P-value |
| --- | --- | --- | --- | --- | --- |
| Membrane Tissue Factor Average Final H Score, median (min,max) | N = 32 | 46.92 (0.67, 231.67) | 20.33 (0.67, 205) | 67.5 (23.67, 231.67) | 0.006 |
| Cytoplasm Tissue Factor Average Final H Score, median (min,max) | N = 32 | 29.16 (2, 180) | 18.33 (2, 123.33) | 61.67 (25, 180) | 0.012 |
| Trop 2 Membrane Average Final H Score, median (min,max) | N = 32 | 106.66 (0, 276.67) | 120.33 (15, 276.67) | 85 (0, 225) | 0.077 |
| Trop 2 Cytoplasm Average Final H Score, median (min,max) | N = 32 | 109.16 (40, 226.67) | 110 (73, 226.67) | 107.5 (40, 145) | 0.274 |
| Nectin 4 Membrane Average Final H Score, median (min,max) | N = 32 | 5.42 (0, 136.67) | 3.33 (0, 136.67) | 7.5 (0, 115) | 0.456 |
| Nectin 4 Cytoplasm Average Final H Score, median (min,max) | N = 32 | 74.16 (0, 171.67) | 88.33 (0, 171.67) | 46.67 (0, 126.67) | 0.131 |

HPV status
| Patient Characteristics |  | Total<br>N=32 | negative N=16 | positive N=16 | P-value |
| --- | --- | --- | --- | --- | --- |
| Membrane Tissue Factor Average Final H Score, median (min,max) | N = 32 | 46.92 (0.67, 231.67) | 69.59 (1.33, 231.67) | 18.75 (0.67, 123.33) | 0.003 |
| Cytoplasm Tissue Factor Average Final H Score, median (min,max) | N = 32 | 29.16 (2, 180) | 59.17 (2.33, 180) | 17.66 (2, 98.33) | 0.007 |
| Trop 2 Membrane Average Final H Score, median (min,max) | N = 32 | 106.66 (0, 276.67) | 96.66 (0, 225) | 107.5 (15, 276.67) | 0.207 |
| Trop 2 Cytoplasm Average Final H Score, median (min,max) | N = 32 | 109.16 (40, 226.67) | 107.91 (40, 145) | 124.16 (85.33, 226.67) | 0.073 |
| Nectin 4 Membrane Average Final H Score, median (min,max) | N = 32 | 5.42 (0, 136.67) | 12.09 (0, 136.67) | 3.33 (0, 80) | 0.618 |
| Nectin 4 Cytoplasm Average Final H Score, median (min,max) | N = 32 | 74.16 (0, 171.67) | 85.83 (0, 171.67) | 65 (0, 168.33) | 0.720 |

Primary Tumor Grade
| Patient Characteristics |  | Total N=32 | 2 N=13 | 3 N=19 | P-value |
| --- | --- | --- | --- | --- | --- |
| Membrane Tissue Factor Average Final H Score, median (min,max) | N = 32 | 46.92 (0.67, 231.67) | 47.5 (4, 205) | 46.33 (0.67, 231.67) | 0.323 |
| Cytoplasm Tissue Factor Average Final H Score, median (min,max) | N = 32 | 29.16 (2, 180) | 28.33 (3, 123.33) | 30 (2, 180) | 0.367 |
| Trop 2 Membrane Average Final H Score, median (min,max) | N = 32 | 106.66 (0, 276.67) | 93.33 (0, 185) | 136.67 (15, 276.67) | 0.053 |
| Trop 2 Cytoplasm Average Final H Score, median (min,max) | N = 32 | 109.16 (40, 226.67) | 106.67 (90, 145) | 110 (40, 226.67) | 0.513 |
| Nectin 4 Membrane Average Final H Score, median (min,max) | N = 32 | 5.42 (0, 136.67) | 15 (0, 69.33) | 3.33 (0, 136.67) | >0.99 |
| Nectin 4 Cytoplasm Average Final H Score, median (min,max) | N = 32 | 74.16 (0, 171.67) | 60 (0, 125) | 83.33 (0, 171.67) | 0.233 |

p16 Status
| Patient Characteristics |  | Total<br>N=32 | negative N=17 | positive N=15 | P-value |
| --- | --- | --- | --- | --- | --- |
| Membrane Tissue Factor Average Final H Score, median (min,max) | N = 32 | 46.92 (0.67, 231.67) | 71.67 (23.67, 231.67) | 15 (0.67, 108.33) | <0.001 |
| Cytoplasm Tissue Factor Average Final H Score, median (min,max) | N = 32 | 29.16 (2, 180) | 61.67 (15.67, 180) | 11.67 (2, 98.33) | <0.001 |
| Trop 2 Membrane Average Final H Score, median (min,max) | N = 32 | 106.66 (0, 276.67) | 85 (0, 225) | 120.33 (15, 276.67) | 0.052 |
| Trop 2 Cytoplasm Average Final H Score, median (min,max) | N = 32 | 109.16 (40, 226.67) | 107.5 (40, 145) | 135 (85.33, 226.67) | 0.041 |
| Nectin 4 Membrane Average Final H Score, median (min,max) | N = 32 | 5.42 (0, 136.67) | 7.5 (0, 115) | 3.33 (0, 136.67) | 0.939 |
| Nectin 4 Cytoplasm Average Final H Score, median (min,max) | N = 32 | 74.16 (0, 171.67) | 70 (0, 126.67) | 78.33 (0, 171.67) | 0.596 |

**Supplemental Table 3.** Descriptive statistics of patient population. Percentage is given in the parentheses.

|  | Total (n=33) | HPV Positive (n=17) | HPV Negative (n=16) | p-value |
| --- | --- | --- | --- | --- |
| Race/Ethnicity (%) |  |  |  | 0.31 |
| Caucasian/White | 20 (60.6) | 11 (64.7) | 9 (56.3) |  |
| African American/Black | 3 (9.1) | 3 (17.7) |  |  |
| Hispanic | 10 (30.3) | 3 (17.7) | 7 (43.8) |  |
| Median Age (years, IQR) | 64 (51.5 – 70) | 66 (54 – 69.5) | 58 (38 – 70) | 0.49 |
| Median BMI (IQR) | 30.7 (22.5 – 48) | 29.4 (28.2 – 38.3) | 32 (26.8 – 37) | 0.98 |
| Median Follow-up (months; IQR) | 19.3 (5.5 – 42.2) | 21.4 (7.2 – 37.3) | 18.0 (5.4 – 56.1) | 0.94 |
| P53 |  |  |  | 0.66 |
| Yes | 25 (75.8) | 14 (82.4) | 11 (68.8) |  |
| No | 8 (18.4) | 3 (17.7) | 5 (31.3) |  |
| LVI (%) |  |  |  | 0.96 |
| Yes | 24 (72.7%) | 12 (70.6) | 12 (75.0) |  |
| No | 9 (27.3) | 5 (29.4) | 4 (25.0) |  |
| PNI (%) |  |  |  | 0.92 |
| Yes | 12 (36.4) | 5 (29.4) | 7 (43.8) |  |
| No | 10 (30.3) | 5 (29.4) | 5 (31.3) |  |
| N/a | 11 (33.3) | 7 (41.2) | 4 (25.0) |  |
| Metastasis (%) |  |  |  | 0.42 |
| Yes | 19 (57.6) | 7 (41.2) | 10 (58.8) |  |
| No | 14 (42.4) | 10 (58.8) | 6 (37.5) |  |
| Tumor grade (%) |  |  |  | 0.98 |
| G2 | 13 (39.4) | 7 (41.2) | 6 (37.5) |  |
| G3 | 20 (60.6) | 10 (58.8) | 10 (58.8) |  |
| Clinical tumor stage (%) |  |  |  | 0.66 |
| T1 | 6 (18.2) | 1 (5.9) | 5 (31.3) |  |
| T2 | 17 (51.5) | 10 (58.8) | 7 (43.8) |  |
| T3 | 8 (24.2) | 5 (29.4) | 3 (18.8) |  |
| T4 | 1 (3.0) |  | 1 (6.3) |  |
| Tx | 1 (3.0) | 1 (5.9) |  |  |
| Clinical nodal stage |  |  |  | 0.33 |
| N0 | 19 (57.6) | 11 (64.7) | 8 (50.0) |  |
| N1 | 9 (27.3) | 2 (11.8) | 7 (43.8) |  |
| N2 | 3 (9.1) | 3 (17.7) |  |  |
| N3 | 1 (3.0) | 1 (5.9) |  |  |
| Nx | 1 (3.0) |  | 1 (6.3) |  |
| Pathologic tumor stage |  |  |  | >0.99 |
| T2 | 23 (69.7) | 12 (70.6) | 11 (68.8) |  |
| T3 | 9 (27.3) | 5 (29.4) | 4 (25.0) |  |
| T4 | 1 (3.0) |  | 1 (6.3) |  |
| Pathologic nodal stage |  |  |  | 0.22 |
| N0 | 12 (36.4) | 10 (58.8) | 2 (12.5) |  |
| N1 | 7 (21.2) | 3 (17.7) | 3 (18.8) |  |
| N2 | 4 (12.1) | 1 (5.9) | 2 (12.5) |  |
| N3 | 8 (24.2) | 2 (11.8) | 8 (50.0) |  |
| Nx | 2 (6.1) | 1 (5.9) | 1 (6.3) |  |
| Primary Procedure (%) |  |  |  | 0.97 |
| Partial penectomy | 22 (66.7) | 11 (64.7) | 11 (68.8) |  |
| Total penectomy | 11 (33.3) | 6 (35.3) | 5 (31.3) |  |
| Histology (%) |  |  |  | 0.41 |
| Usual | 28 (84.9) | 13 (76.5) | 15 (93.8) |  |
| Basaloid | 5 (15.2) | 4 (23.5) | 1 (6.3) |  |
| Recurrence (%) |  |  |  | 0.01 |
| Yes | 9 (27.3) | 4 (23.5) | 11 (68.8) |  |
| No | 24 (72.7) | 13 (76.5) | 5 (31.3) |  |
| Survival (%) |  |  |  | 0.30 |
| Alive | 17 (51.5) | 11 (64.7) | 6 (37.5) |  |
| Dead | 16 (48.5) | 6 (35.3) | 10 (58.8) |  |

## References

1. Pow-Sang, M. R., Ferreira, U., Pow-Sang, J. M., Nardi, A. C. and Destefano, V. Epidemiology and natural history of penile cancer Urology. 2010; 76: S2–6.

2. Misra, S., Chaturvedi, A. and Misra, N. C. Penile carcinoma: a challenge for the developing world Lancet Oncol. 2004; 5: 240–247.

3. Pettaway, C. A., Pagliaro, L., Theodore, C. and Haas, G. Treatment of visceral, unresectable, or bulky/unresectable regional metastases of penile cancer Urology. 2010; 76: S58–65.

4. Vreeburg, M. T. A., Donswijk, M. L., Albersen, M., Parnham, A., Ayres, B., Protzel, C., Pettaway, C., Spiess, P. E. and Brouwer, O. R. New EAU/ASCO guideline recommendations on sentinel node biopsy for penile cancer and remaining challenges from a nuclear medicine perspective Eur J Nucl Med Mol Imaging. 2024; 51: 2861–2868.

5. Pagliaro, L. C., Williams, D. L., Daliani, D., Williams, M. B., Osai, W., Kincaid, M., Wen, S., Thall, P. F. and Pettaway, C. A. Neoadjuvant paclitaxel, ifosfamide, and cisplatin chemotherapy for metastatic penile cancer: a phase II study J Clin Oncol. 2010; 28: 3851–3857.

6. Ottenhof, S. R., de Vries, H. M., Doodeman, B., Vrijenhoek, G. L., van der Noort, V., Donswijk, M. L., de Feijter, J. M., Schaake, E. E., Horenblas, S., Brouwer, O. R., van der Heijden, M. S. and Pos, F. J. A Prospective Study of Chemoradiotherapy as Primary Treatment in Patients With Locoregionally Advanced Penile Carcinoma Int J Radiat Oncol Biol Phys. 2023; 117: 139–147.

7. Brouwer, O. R., Albersen, M., Parnham, A., Protzel, C., Pettaway, C. A., Ayres, B., Antunes-Lopes, T., Barreto, L., Campi, R., Crook, J., Fernandez-Pello, S., Greco, I., van der Heijden, M. S., Johnstone, P. A. S., Kailavasan, M., Manzie, K., Marcus, J. D., Necchi, A., Oliveira, P., Osborne, J., Pagliaro, L. C., Garcia-Perdomo, H. A., Rumble, R. B., Sachdeva, A., Sakalis, V. I., Zapala, L., Sanchez Martinez, D. F., Spiess, P. E. and Tagawa, S. T. European Association of Urology-American Society of Clinical Oncology Collaborative Guideline on Penile Cancer: 2023 Update Eur Urol. 2023; 83: 548-560.

8. Rose, K. M., Pham, R., Zacharias, N. M., Ionescu, F., Paravathaneni, M., Marchetti, K. A., Sanchez, D., Mustasam, A., Sandstrom, R., Vikram, R., Dhillon, J., Rao, P., Schneider, A., Pagliaro, L., Alifrangis, C., Albersen, M., Roussel, E., Master, V. A., Nazha, B., Hernandez, C., Moses, K. A., Protzel, C., Montgomery, J., Angel, M., Tobias-Machado, M., Spiess, P. E., Pettaway, C. A. and Chahoud, J. Neoadjuvant platinum-based chemotherapy and lymphadenectomy for penile cancer: an international, multi-institutional, real-world study J Natl Cancer Inst. 2024; 116: 966–973.

9. Wang, J., Pettaway, C. A. and Pagliaro, L. C. Treatment for Metastatic Penile Cancer After First-line Chemotherapy Failure: Analysis of Response and Survival Outcomes Urology. 2015; 85: 1104–1110.

10. Pagliaro, L. C., Tekin, B., Gupta, S. and Herrera Hernandez, L. Therapeutic Targets in Advanced Penile Cancer: From Bench to Bedside Cancers (Basel). 2024; 16:

11. Grass, G. D., Chahoud, J., Lopez, A., Dhillon, J., Eschrich, S. A., Johnstone, P. A. S. and Spiess, P. E. An Analysis of Nectin-4 (PVRL4) in Penile Squamous Cell Carcinoma Eur Urol Open Sci. 2023; 49: 1-5.

12. Tekin, B., Cheville, J. C., Herrera Hernandez, L., Negron, V., Smith, C. Y., Jenkins, S. M., Dasari, S., Enninga, E. A. L., Norgan, A. P., Menon, S., Cubilla, A. L., Whaley, R. D., Jimenez, R. E., Thompson, R. H., Leibovich, B. C., Karnes, R. J., Boorjian, S. A., Pagliaro, L. C., Erickson, L. A., Guo, R. and Gupta, S. Assessment of PD-L1, TROP2, and nectin-4 expression in penile squamous cell carcinoma Hum Pathol. 2023; 142: 42-50.

13. Versteeg, H. H. Tissue Factor: Old and New Links with Cancer Biology Semin Thromb Hemost. 2015; 41: 747–755.

14. Rondon, A. M. R., Kroone, C., Kapteijn, M. Y., Versteeg, H. H. and Buijs, J. T. Role of Tissue Factor in Tumor Progression and Cancer-Associated Thrombosis Semin Thromb Hemost. 2019; 45: 396–412.

15. Coleman, R. L., Lorusso, D., Gennigens, C., Gonzalez-Martin, A., Randall, L., Cibula, D., Lund, B., Woelber, L., Pignata, S., Forget, F., Redondo, A., Vindelov, S. D., Chen, M., Harris, J. R., Smith, M., Nicacio, L. V., Teng, M. S. L., Laenen, A., Rangwala, R., Manso, L., Mirza, M., Monk, B. J., Vergote, I. and innova, T. V. G. O. G. E.-c. C. Efficacy and safety of tisotumab vedotin in previously treated recurrent or metastatic cervical cancer (innovaTV 204/GOG-3023/ENGOT-cx6): a multicentre, open-label, single-arm, phase 2 study Lancet Oncol. 2021; 22: 609-619.

16. Vergote, I., Gonzalez-Martin, A., Fujiwara, K., Kalbacher, E., Bagameri, A., Ghamande, S., Lee, J. Y., Banerjee, S., Maluf, F. C., Lorusso, D., Yonemori, K., Van Nieuwenhuysen, E., Manso, L., Woelber, L., Westermann, A., Covens, A., Hasegawa, K., Kim, B. G., Raimondo, M., Bjurberg, M., Cruz, F. M., Angelergues, A., Cibula, D., Barraclough, L., Oaknin, A., Gennigens, C., Nicacio, L., Teng, M. S. L., Whalley, E., Soumaoro, I., Slomovitz, B. M. and innova, T. V. E.-c. G. O. G. C. Tisotumab Vedotin as Second-or Third-Line Therapy for Recurrent Cervical Cancer N Engl J Med. 2024; 391: 44–55.

17. Chahoud, J., Zacharias, N. M., Pham, R., Qiao, W., Guo, M., Lu, X., Alaniz, A., Segarra, L., Martinez-Ferrer, M., Gleber-Netto, F. O., Pickering, C. R., Rao, P. and Pettaway, C. A. Prognostic Significance of p16 and Its Relationship with Human Papillomavirus Status in Patients with Penile Squamous Cell Carcinoma: Results of 5 Years Follow-Up Cancers (Basel). 2022; 14: 6024.

18. Jawhar, N. M. Tissue Microarray: A rapidly evolving diagnostic and research tool Ann Saudi Med. 2009; 29: 123–127.

19. Tessier-Cloutier, B., Kortekaas, K. E., Thompson, E., Pors, J., Chen, J., Ho, J., Prentice, L. M., McConechy, M. K., Chow, C., Proctor, L., McAlpine, J. N., Huntsman, D. G., Gilks, C. B., Bosse, T. and Hoang, L. N. Major p53 immunohistochemical patterns in in situ and invasive squamous cell carcinomas of the vulva and correlation with TP53 mutation status Mod Pathol. 2020; 33: 1595–1605.

20. Kobel, M. and Kang, E. Y. The Many Uses of p53 Immunohistochemistry in Gynecological Pathology: Proceedings of the ISGyP Companion Society Session at the 2020 USCAP Annual9 Meeting Int J Gynecol Pathol. 2021; 40: 32–40.

21. Trias, I., Saco, A., Marimon, L., Lopez Del Campo, R., Manzotti, C., Ordi, O., Del Pino, M., Perez, F. M., Vega, N., Alos, S., Martinez, A., Rodriguez-Carunchio, L., Reig, O., Jares, P., Teixido, C., Ajami, T., Corral-Molina, J. M., Algaba, F., Ribal, M. J., Ribera-Cortada, I. and Rakislova, N. P53 in Penile Squamous Cell Carcinoma: A Pattern-Based Immunohistochemical Framework with Molecular Correlation Cancers (Basel). 2023; 15:

22. Kashofer, K., Winter, E., Halbwedl, I., Thueringer, A., Kreiner, M., Sauer, S. and Regauer, S. HPV-negative penile squamous cell carcinoma: disruptive mutations in the TP53 gene are common Mod Pathol. 2017; 30: 1013–1020.

23. Chahoud, J., Gleber-Netto, F. O., McCormick, B. Z., Rao, P., Lu, X., Guo, M., Morgan, M. B., Chu, R. A., Martinez-Ferrer, M., Eterovic, A. K., Pickering, C. R. and Pettaway, C. A. Whole-exome Sequencing in Penile Squamous Cell Carcinoma Uncovers Novel Prognostic Categorization and Drug Targets Similar to Head and Neck Squamous Cell Carcinoma Clin Cancer Res. 2021; 27: 2560–2570.

24. de Bono, J. S., Harris, J. R., Burm, S. M., Vanderstichele, A., Houtkamp, M. A., Aarass, S., Riisnaes, R., Figueiredo, I., Nava Rodrigues, D., Christova, R., Olbrecht, S., Niessen, H. W. M., Ruuls, S. R., Schuurhuis, D. H., Lammerts van Bueren, J. J., Breij, E. C. W. and Vergote, I. Systematic study of tissue factor expression in solid tumors Cancer Rep (Hoboken). 2023; 6: e1699.

25. Zhao, X. T., Cheng, C., Gou, J. H., Yi, T., Qian, Y. P., Du, X. and Zhao, X. Expression of tissue factor in human cervical carcinoma tissue Experimental and Therapeutic Medicine. 2018; 16: 4075–4081.

26. Randall, L. M., Walker, A. J., Jia, A. Y., Miller, D. T. and Zamarin, D. Expanding Our Impact in Cervical Cancer Treatment: Novel Immunotherapies, Radiation Innovations, and Consideration of Rare Histologies Am Soc Clin Oncol Educ Book. 2021; 41: 252–263.

27. Kasthuri, R. S., Taubman, M. B. and Mackman, N. Role of Tissue Factor in Cancer Journal of Clinical Oncology. 2009; 27: 4834–4838.

28. Sawada, M., Miyake, S., Ohdama, S., Matsubara, O., Masuda, S., Yakumaru, K. and Yoshizawa, Y. Expression of tissue factor in non-small-cell lung cancers and its relationship to metastasis British Journal of Cancer. 1999; 79: 472–477.

29. Ueno, T., Toi, M., Koike, M., Nakamura, S. and Tominaga, T. Tissue factor expression in breast cancer tissues: its correlation with prognosis and plasma concentration British Journal of Cancer. 2000; 83: 164–170.

30. Shigemori, C., Wada, H., Matsumoto, K., Shiku, H., Nakamura, S. and Suzuki, H. Tissue factor expression and metastatic potential of colorectal cancer Thrombosis and Haemostasis. 1998; 80: 894–898.

31. Seto, S., Onodera, H., Kaido, T., Yoshikawa, A., Ishigami, S., Arii, S. and Imamura, M. Tissue factor expression in human colorectal carcinoma - Correlation with hepatic metastasis and impact on prognosis Cancer. 2000; 88: 295–301.

32. Elst, L., Philips, G., Vandermaesen, K., Bassez, A., Lodi, F., Vreeburg, M. T. A., Brouwer, O. R., Schepers, R., Van Brussel, T., Mohanty, S. K., Parwani, A. V., Spans, L., Vanden Bempt, I., Jacomen, G., Baldewijns, M., Lambrechts, D. and Albersen, M. Single-cell Atlas of Penile Cancer Reveals TP53 Mutations as a Driver of an Aggressive Phenotype, Irrespective of Human Papillomavirus Status, and Provides Clues for Treatment Personalization Eur Urol. 2024; 86: 114–127.

33. Breij, E. C. W., de Goeij, B. E. C. G., Verploegen, S., Schuurhuis, D. H., Amirkhosravi, A., Francis, J., Miller, V. B., Houtkamp, M., Bleeker, W. K., Satijn, D. and Parren, P. W. H. I. An Antibody-Drug Conjugate That Targets Tissue Factor Exhibits Potent Therapeutic Activity against a Broad Range of Solid Tumors Cancer Research. 2014; 74: 1214–1226.

